# DNA framework array enables ultra-high throughput DNA synthesis

**DOI:** 10.1101/2025.05.30.657018

**Authors:** Chunhong Li, Yishakejiang Saimaiti, Min Li, Shaopeng Wang, Fangfei Yin, Zheng Fang, Sisi Jia, Fei Wang, Dekai Ye, Lihua Wang, Xiaolei Zuo, Chunhai Fan

## Abstract

High-throughput DNA synthesis has revolutionized synthetic biology, molecular diagnostics, genome engineering, and DNA data storage by enabling the scalable and precise construction of nucleotide-based information systems. However, existing high-throughput synthesis technologies rely on top-down fabrication strategies to maximize throughput, yet they are inherently constrained by physical limitations. These constraints make it exceedingly difficult to achieve single-molecule level synthesis control that is essential for advancing next-generation DNA synthesis. Here, we present a DNA framework-based bottom-up enzymatic synthesis strategy that enables single-molecule level control of DNA synthesis with high-throughput. Leveraging the nanoscale addressability of DNA frameworks, we achieved a synthesis site pitch of 10.9 nm—183-fold smaller than that of previously reported electrode arrays. Theoretical projections suggest that this approach could scale synthesis throughput to 5.3 × 10^11^ sequences per square centimeter, representing a four-order-of-magnitude improvement over existing synthesis arrays. This approach lays the foundation for gigabyte-scale DNA writing and offers a highly accessible pathway toward large-scale DNA synthesis for emerging applications such as molecular data storage.

## Introduction

High-throughput DNA synthesis has transformed the ability to generate diverse and precise nucleotide sequences at an unprecedented scale, driving breakthroughs in many fields such as synthetic biology, molecular diagnostics, and genome engineering.^1^ By enabling the rapid and cost-effective assembly of genetic constructs, high-throughput synthesis accelerates the design-build-test cycle for engineered biological systems, facilitates the development of novel therapeutics, and enhances our understanding of genetic functions^.1,2^ Among its many applications, DNA data storage has emerged as a particularly promising solution to address the growing global data crisis. With global data projected to reach 1000 zettabytes by 2035,^3^ conventional storage media are facing fundamental limitations in scalability, energy consumption, and longevity.^4^ In contrast, DNA, with an anticipated practical storage density exceeding 60 petabytes per cubic centimeter, offers an ultra-high-density and durable alternative.^5-9^ However, a major bottleneck remains in the efficiency and scalability of DNA synthesis, as the current throughput for writing digital data into DNA is insufficient to meet the demands of large-scale storage applications.^10^

To enhance throughput, several innovative approaches to DNA synthesis have been developed, including inkjet printing-based synthesis, photomask array-based synthesis, and electrode array-based synthesis.^11^ Inkjet printing technology has demonstrated the ability to synthesize tens of thousands of DNA sequences per square centimeter using standard phosphoramidite chemistry.^12^ More recently, the integration of enzymatic DNA synthesis with inkjet printing has achieved a synthesis spot density of up to 10,560 on a 75 mm × 25 mm glass slide.^13^ The rapid advances in lithography have further reduced the spacing between synthesis sites, enabling top-down photolithographic DNA synthesis technology to parallelly synthesize hundreds of thousands of DNA strands per square centimeter (Fig. 1a).^14^ Notably, Microsoft and the University of Washington recently developed a sub-millimeter DNA synthesis chip utilizing a micro-nanoelectrode array.^15^ Through a 130 nm photolithography process, the size of the synthesis sites and pitch were reduced to 650 nm and 2 μm respectively, enabling a DNA synthesis throughput of 25 million oligonucleotides per square centimeter. This enhanced synthesis throughput has resulted in write throughputs being promoted to megabytes per second. Despite the significant advancement, the current writing throughput remains insufficient for practical data storage applications. More critically, the scalability of these techniques is fundamentally constrained by their reliance on top-down nanofabrication. The need for high-precision photolithography, microfluidic patterning, and electrode arrays imposes substantial cost and infrastructure requirements, limiting accessibility for smaller laboratories.^16-18^ Additionally, intrinsic physical barriers—including droplet size, micromirror resolution, and light diffraction effects—further restrict the miniaturization and throughput potential of these methods. Overcoming these barriers necessitates the development of novel synthesis paradigms that move beyond traditional top-down fabrication, enabling more accessible and scalable DNA synthesis technologies.

**Fig. 1.**
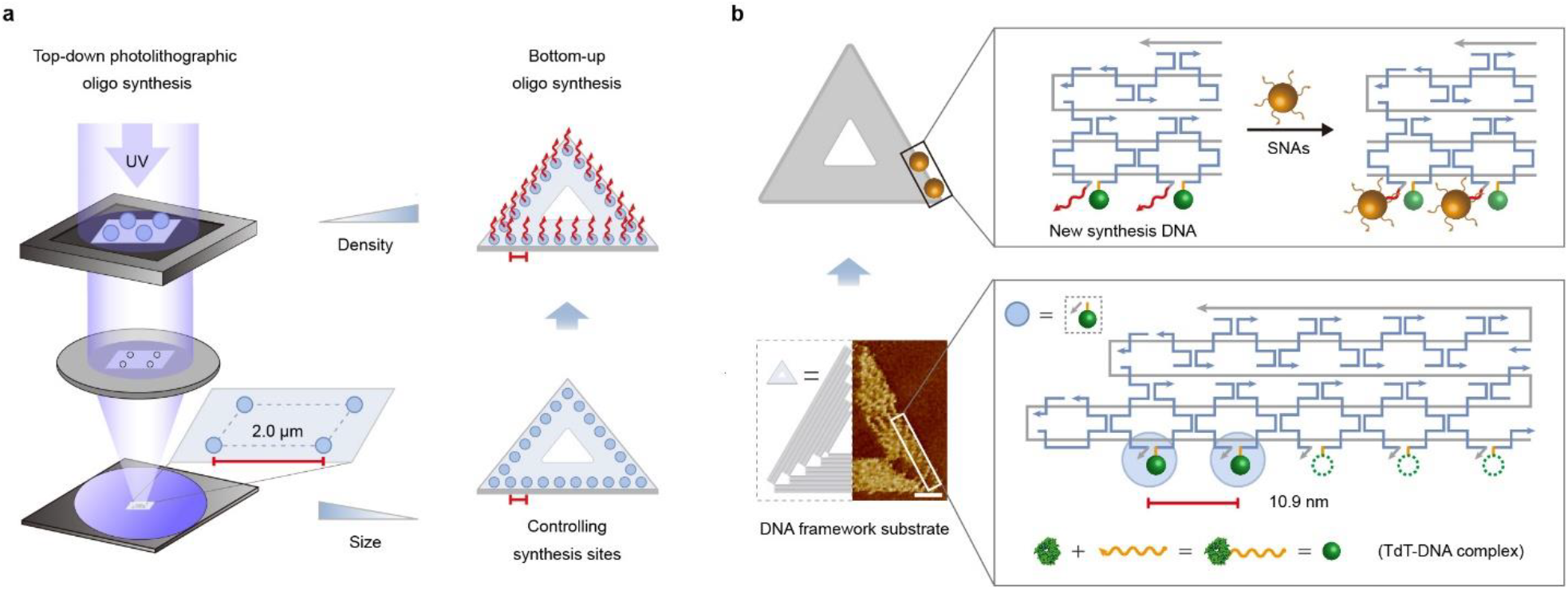
The bottom-up high-throughput enzymatic DNA synthesis based on DNA framework. (a) schematic illustration of the top-down photolithographic oligo synthesis and bottom-up enzymatic oligo synthesis based on DNA framework. In the photolithography process, the photosensitive polymer forms a mask on the chip for etching. The minimum size of the pattern is determined by the wavelength of the light used. In the bottom-up oligo synthesis method, the position and distance of the synthesis site are regulated by the DNA framework. (b) Scheme illustrating the precise regulation of synthesis site distance by DNA framework.

DNA nanotechnology, a prototypical bottom-up approach, has garnered significant attention in the realm of nanofabrication and been used to produce framework structures from nanoscale to microscale.^19^ The inherent bottom-up nature of DNA nanotechnology ensures the assembly of framework structures with well-defined spacing, orientations and stereo-relationships. Moreover, the nanometer-scale addressability of DNA frameworks enables the precise organization of a variety of functional molecules, including proteins, nucleic acids, inorganic nanoparticles, and organic dyes.^20-23^ Here we report the development of DNA framework-based high throughput enzymatic DNA synthesis strategy (Fig. 1). Leveraging the structural rigidity and nanometer-scale addressability of DNA framework array, we precisely position single-stranded DNA conjugated TdT, controlling synthesis sites with nanometer precision in terms of number, orientation, and spacing. This approach reduces the pitch between synthesis size to 10.9 nm, leading to a synthesis throughput of 5.3×10 ^11^ sequences per square centimeter—a four-order-of-magnitude improvement over current methods. Using this method, we successfully encoded the acronym “Next Generation Data Store (NGDS)” into homopolymeric DNA extensions synthesized enzymatically. Furthermore, this framework-based approach enables single-molecule-level control of DNA synthesis at designated sites. In contrast to traditional top-down manufacturing, which has historically adhered to Moore’s Law but faces challenges in sustainability for the future, our DNA framework-based approach achieves unprecedented spatial precision, with a feature pitch of 10.9 nm—surpassing the current transistor gate length (18 nm)^24-27^ and exceeding the transistor density of silicon-based electronic memory by an order of magnitude.^28,29^

## Results and Discussion Controlling the synthesis sites

In recent years, TdT-based enzymatic DNA synthesis has attracted significant attention for its ability to generate longer DNA strands with greater accuracy, faster synthesis rates, and fewer byproducts compared to traditional chemical synthesis methods. Notably, the protein nature of TdT enables seamless integration with DNA nanotechnology, facilitating straightforward modification and assembly within DNA frameworks. To precisely regulate the synthesis sites on the DNA framework, TdT was covalently conjugated to a linker DNA, which serves as an anchor to hybridize with capture strands displayed on the surface of DNA framework. The TdT-DNA complex was prepared using the heterobifunctional crosslinker succinimidyl 3-(2-pyridyldithio) propionate (SPDP) (Fig. 2a). The successful preparation of the complex was confirmed by PAGE analysis, which revealed that the TdT-DNA complex exhibited much slower migration than the linker DNA alone (Fig. 2b). Absorption spectroscopy determined that approximately two linker DNA molecules were labeled on each TdT.

**Figure 2.**
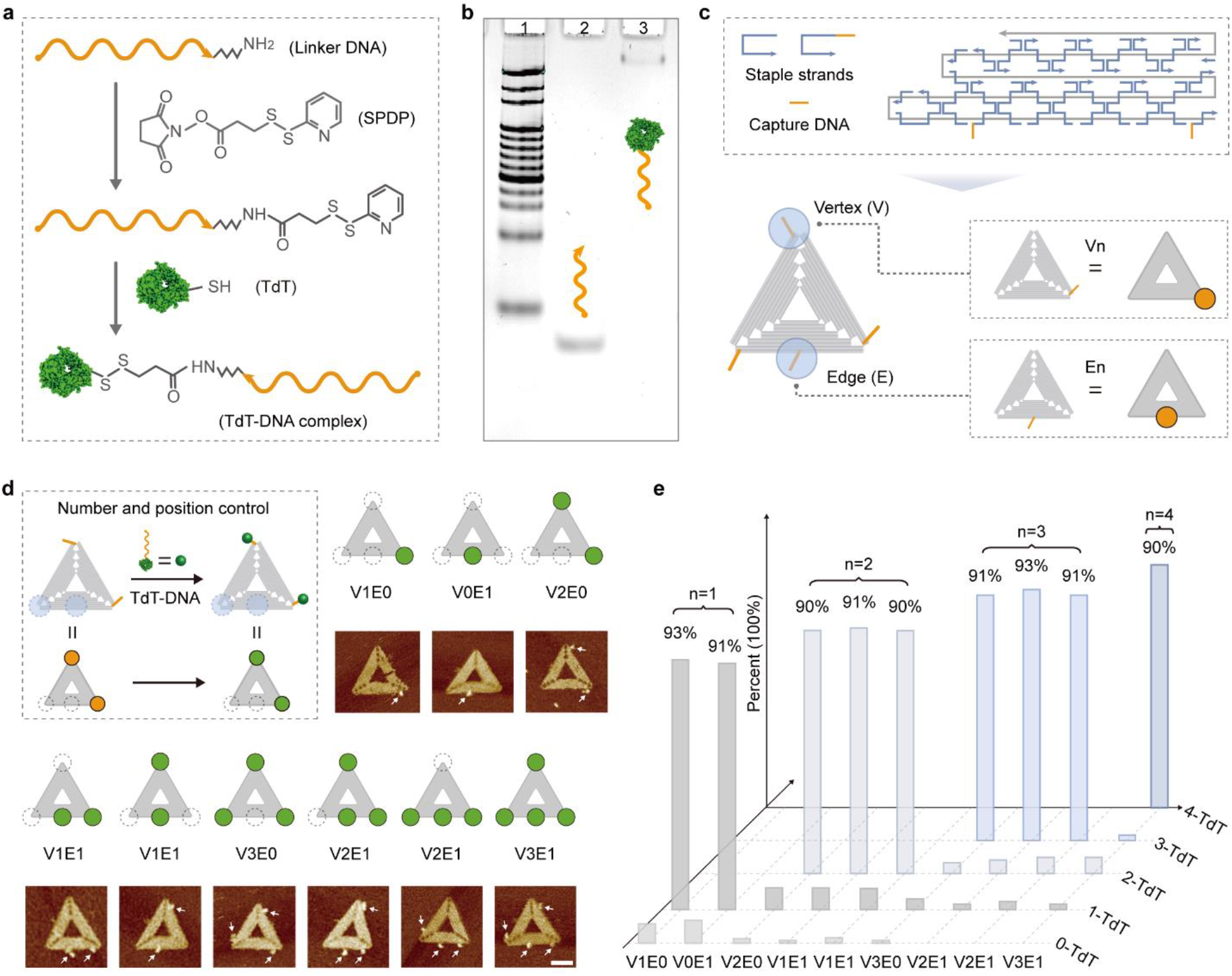
Number and position control of synthesis sites. (a) Schematic diagram of the preparation of TdT-DNA complex by heterobifunctional crosslinker succinimidyl 3-(2-pyridyldithio) propionate (SPDP). The SPDP was used to cross link the amidogen-modified oligonucleotide to the cysteine of TdT. (b) PAGE characterization of TdT-DNA complex. (c) Schematic diagram of the TdT capture chain designed at special sites on the interface of DNA framework. We selected three vertices (V) and one midpoint of an edge (E) of the DNA framework as feature synthetic sites. (d) Scheme of TdT-DNA complex assembly with DNA framework and the AFM characterization of different number of TdT-DNA complexes assembly with DNA framework at different position. Scale bar, 50 nm. (e) Efficiency statistics of TdT-DNA complex assembly with DNA framework.

After successful conjugation, the TdT-DNA complex was assembled onto DNA framework via hybridization with the capture strands. We chose triangular DNA origami, one of the DNA frameworks with precisely customizable shape and size, as the scaffold for TdT-DNA complex assembly. A single scaffold DNA framework can theoretically provide hundreds of globally unique addresses, allowing for the spatially addressable arrangement of synthesis sites with sub-nanometer resolution. To demonstrate precise control over synthesis sites, we strategically designed distinct synthesis locations at the vertices and midpoints of the DNA framework to program the number and position of TdT molecules accurately (Fig. 2c). The successful assembly of 1-4 TdT molecules at specific positions on DNA framework was validated using AFM imaging (Fig. 2d), with over 90% correct assembly yield for each design (Fig. 2e), indicating the accurate regulation of synthesis sites.

### Enzymatic DNA synthesis at single-molecule level

To achieve precise control over the TdT enzymatic DNA synthesis process, it is crucial to understand the detailed mechanisms and dynamics of the reaction. In ultra-high-throughput synthesis, where synthesis sites are confined to the substrate surface and potentially reduced to nanoscale dimensions, understanding the reaction at the single-molecule level on a solid support becomes essential. However, much of our current understanding of TdT-based enzymatic DNA synthesis comes from ensemble-averaged methods that interrogate ensembles of DNA molecules undergoing various stages of the reaction concurrently.

To gain insight into the reaction mechanism at the single-molecule level, we first employed total internal reflection fluorescence microscopy (TIRFM) to examine the transition states and reaction trajectories in TdT-based DNA synthesis, which were transduced into fluorescent signals. In our single-molecule experiments, Alexa fluor 488 modified DNA primers were immobilized on a confocal dish surface in 1′ TdT reaction buffer containing cyanine-3 (Cy3) modified deoxyadenosine triphosphate (dATPs) and TdT (Fig. 3a). The synthesis process was monitored by recording changes in Cy3 fluorescence intensity at DNA primer sites. During TdT-catalyzed DNA synthesis, deoxynucleotide triphosphates (dNTPs) transiently or irreversibly dock with the primer, leading to the incorporation of Cy3 modified dATPs at the 3’ end of Alexa fluor 488 modified bar, 500 nm.

**Figure 3.**
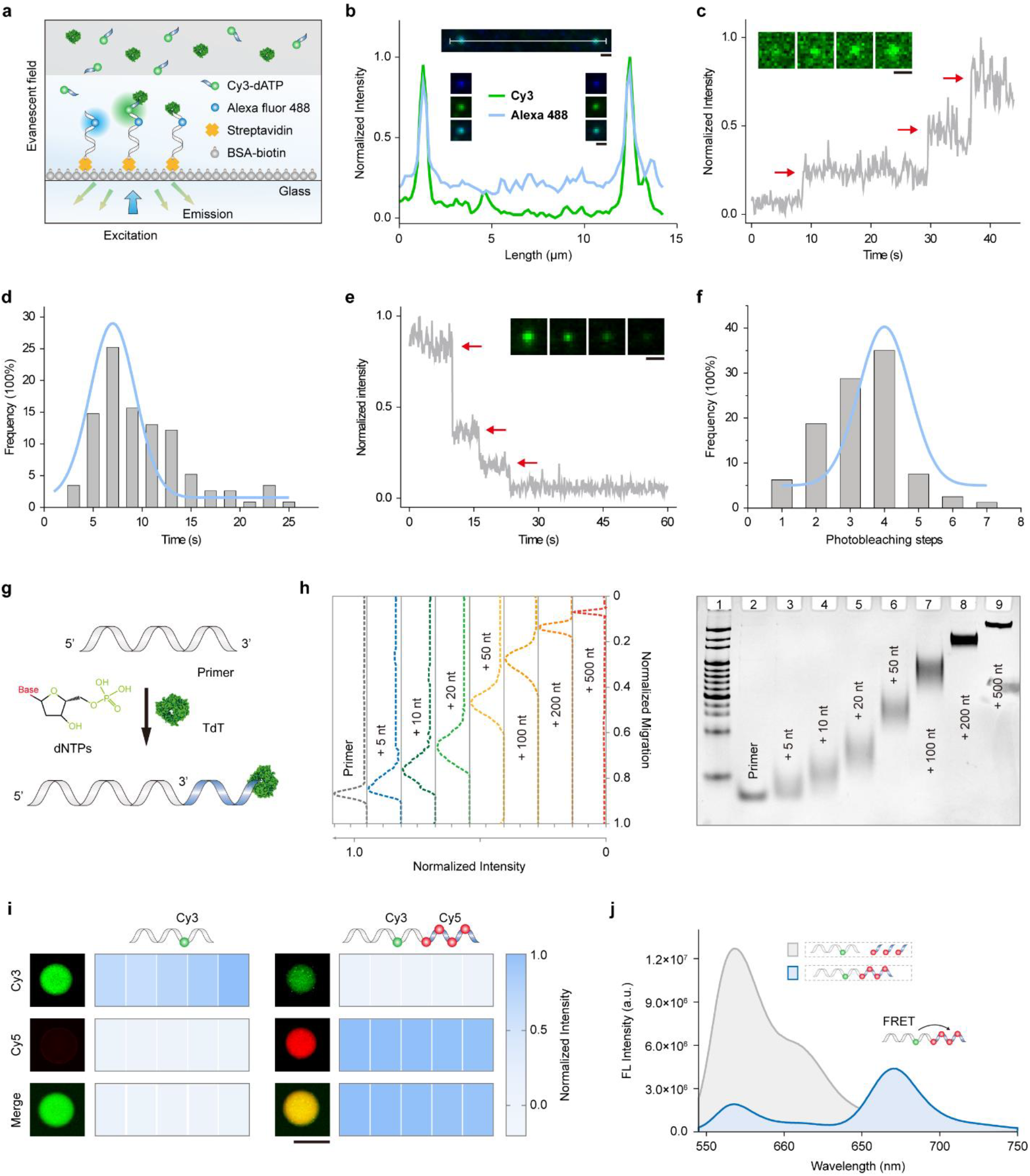
Characterization of TdT-mediated enzymatic DNA synthesis. (a) Schematic representation of TdT-catalyzed DNA synthesis on a glass interface visualized by TIRFM. The glass surface was modified with BSA-biotin, streptavidin, and Alexa Fluor 488 and biotin double-labeled primers. The incorporated nucleotide was Cy3-labeled dATP. (b) Fluorescence intensity profiles along the white line on TIRF images. Insets show corresponding TIRF images, scale bar, 500 nm. (c) Representative intensity–time trajectories for Cy3-labeled dATP incorporation into Alexa Fluor 488-labeled primers, alongside snapshots of raw microscopy images. Scale (d) Distribution of incorporation time for single Cy3-labeled dATP into Alexa Fluor 488-labeled primers. Curves represent Gaussian fits of the distributions (N=115). (e) Representative intensity–time trajectories for Cy3 under TIRFM excitation, showing photobleaching in three steps. Fluorescence dot images for each time point are shown as insets. Scale bar, 500 nm. (f) Statistical distribution of photobleaching steps on TIRF images, with a 4:1 ratio of dNTP to primer. Curves represent Gaussian fits of the distributions (N=80). (g) Schematic illustration of TdT catalyzed dNTP incorporation at the 3’ end of the primer chain. (h) PAGE analysis of TdT-mediated enzymatic DNA synthesis efficiency, with lanes 3-9 showing dTTP-to-primer ratios of 5, 10, 20, 50, 100, 200, and 500, respectively. (i) Fluorescence mapping (left) and intensity profiles (right) of primer incorporation with dNTPs, with or without TdT treatment. The primer was Cy3-labeled, and the incorporated nucleotide was Cy5-labeled dCTP. Reactions were purified by ultrafiltration prior to analysis. Scale bar, 2 mm. (j) Fluorescence spectra of Cy3-labeled primers and free Cy5-labeled dCTP, with or without TdT treatment. primers and an increase in Cy3 fluorescence intensity (Fig. 3a). The perfect colocalization of Alexa fluor 488 (blue) and Cy3 (green) fluorescence indicated successful chain extension (Fig. 3b). Notably, the assay developed here provides precise temporal resolution of TdT reaction dynamics at the single-molecule scale. Fluorescence intensity time trace analysis at individual primer sites revealed a stepwise increase in Cy3 fluorescence, with each step representing a distinct reaction event. The multiple steps observed in the recording time demonstrate successive rounds of Cy3 modified dATPs incorporation at each site (Fig. 3c). The duration of each step correlates directly with the time required for a single nucleotide extension event, statistically calculated to be 9.9 s per nucleotide (Fig. 3d).

Next, we quantified the efficiency of TdT-catalyzed DNA synthesis using single-molecule fluorescence photobleaching, where each step corresponds to the loss of fluorescence from an individual molecule. As expected, the fluorescence intensity of Alexa fluor 488 and Cy3 decreased incrementally over time. The single-step photobleaching of Alexa fluor 488 was consistent with a single modification of the primer (Fig. 3e). For Cy3, the photobleaching curves exhibited stepwise decreases in intensity, ranging from 1 to 7 steps under a 4:1 dNTPs-to-primer ratio, with a 4-step decrease being the most frequent (Fig. 3f). Those findings demonstrate the high efficiency of enzymatic DNA synthesis and suggest that the length of TdT-mediated DNA synthesis can be controlled by adjusting the concentration of dNTPs.

The primary activity of TdT is to indiscriminately add dNTPs to the 3’OH of single-stranded DNA (ssDNA) primer in the absence of a template strand (Fig. 3g). To optimize the conditions of TdT-catalyzed DNA synthesis, we conducted bulk assay to analyze the effects of dNTP type, enzyme concentration, and reaction time on incorporation efficiency. These results allowed us to control and estimate the length of the synthesized homopolymer chain, achieving DNA sequences exceeding 500 nt within 30 min by increasing the dTTP concentration (Fig. 3h). Additionally, we confirmed the incorporation of unnatural nucleotides (fluorescently modified) by TdT in bulk assay. The notable fluorescent intensity and FRET signals observed after TdT-catalyzed incorporation (Fig. 3i, j) indicated successful and efficient chain extension. Taken together, the single-molecule and bulk assay results provided detailed characterization and quantitative insights into the efficiency of TdT-mediated long-chain DNA synthesis.

### Addressable enzymatic DNA synthesis and digital information storage

To achieve high-density enzymatic DNA synthesis, the most space-efficient way is to precisely arrange synthesis sites and minimize the distances between them. Building on previous data indicating that primer extension length can be regulated by dNTP concentration, we reasoned that TdT immobilized on a DNA framework could catalyze primer extension with predictable lengths for primers in its vicinity when provided with appropriate dNTP concentrations (Fig. 4a). The enzymatic activity of TdT-DNA complex was first tested, and it retained robust activity with over 90% of the activity of unmodified TdT (Fig. 4b).

**Figure 4.**
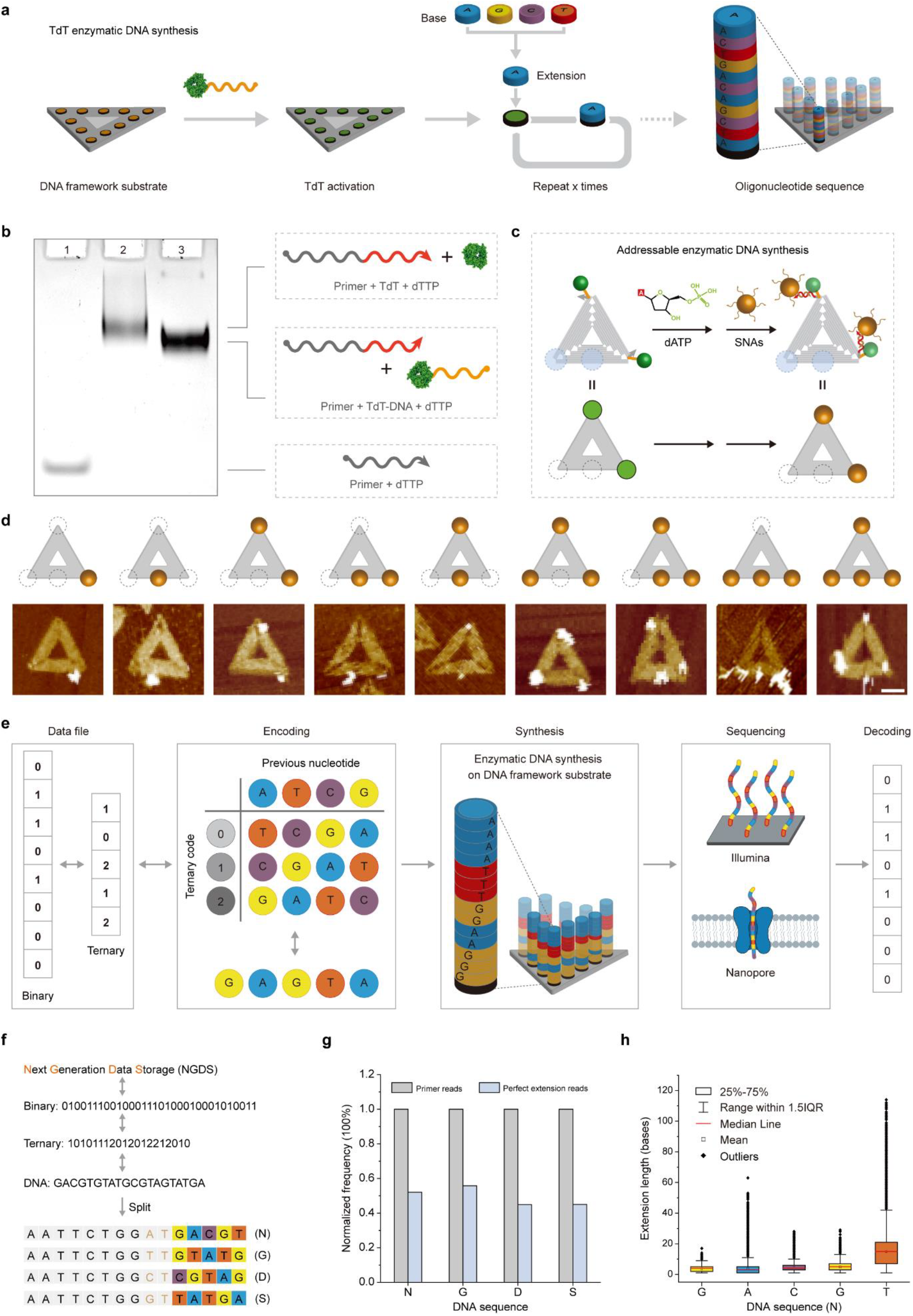
Controlled enzymatic DNA synthesis on the DNA framework interface. (a) Schematic representation of de novo enzymatic DNA synthesis on a DNA framework interface. (b) PAGE analysis of the enzymatic activity of the TdT-DNA complex. (c) Schematic of addressable enzymatic DNA synthesis, with newly synthesized DNA sequences hybridized to SNAs on the DNA framework substrate. (d) AFM imaging of controlled enzymatic DNA synthesis on the DNA framework interface. Scale bar, 50 nm. (e) Schematic of enzymatic DNA synthesis and digital data storage on DNA framework substrate. (f) Schematic of the encoding and mapping process for “Next Generation Data Storage (NGDS)” to DNA sequences. (g) Percentage of perfectly extended reads relative to the total reads containing primers. A perfect extension is defined as a DNA strand that incorporates all five nucleotide bases at the 3’ end of the primer. (h) Distribution of extension lengths for each nucleotide.

To demonstrate controlled and addressable enzymatic DNA synthesis on the DNA framework, poly-A homopolymer sequences were synthesized by adding dATP to the system. The presence of poly-A sequences at the primer ends was characterized using both AFM imaging (Fig. 4c) and fluorescence spectroscopy analysis. Successful poly-A extension on the framework would result in the anchoring of AuNPs densely coated with poly-T oligonucleotide chains (spherical nucleic acids, SNAs) or Cy5-modified poly-T oligonucleotide chains (Fig. 4c). The precise localization of AuNPs at the designated positions on the DNA framework (Fig. 4d) and the significant FRET signal between Cy5 and Cy3 on the primer demonstrate successful and addressable enzymatic DNA synthesis.

Next, we conducted digital data storage by utilizing this addressable enzymatic DNA synthesis platform for data writing (Fig. 4e). User-defined information was first converted to ternary code and then encoded into homopolymeric blocks with unique base transitions using the Huffman coding scheme.^30^ Subsequently, in situ de novo enzymatic DNA synthesis was performed on the DNA framework to write the digital data into DNA. The synthesized DNA strands were then released from the DNA framework through enzymatic cleavage and collected for preservation. To read the data, the DNA strands were sequenced using Illumina sequencing platform. The filtered sequence data was decoded to recover the original information.

To demonstrate the feasibility of this storage strategy, we encoded and synthesized the acronym information for “Next Generation Data Storage (NGDS)”, a message containing 32-bits of ASCII data (Fig. 4f). The information was mapped to a 20-nucleotide sequence, which was further split into four individual sequences. We synthesized all four sequences in parallel and released them from the DNA framework using the endonuclease EcoR I. The success synthesis of all four strands was first confirmed by PAGE analysis. Subsequently, Illumina sequencing was performed to read the synthesized strands. Further analysis revealed that approximately half of the extended sequences perfectly matched the encoded sequence (Fig. 4g). Moreover, the extension length of each nucleotide was generally consistent with our design across all four strands, despite some variations (Fig. 4h). Collectively, these data demonstrate that the addressable enzymatic DNA synthesis platform developed here can be effectively used for data writing in DNA data storage.

### Programming the pitches

The density of DNA synthesis sites is closely related to synthesis throughput, which in turn determines the data writing rate in DNA data storage. This density is inversely proportional to the distance between adjacent synthesis sites (Fig. 5a). Therefore, reducing the distance between synthesis sites would dramatically increase DNA synthesis throughput, thereby enhancing the writing efficiency of DNA data storage. Harnessing the precise nanometer-scale addressability offered by DNA frameworks, we aimed to program the distances between the synthesis sites. We designed a series of synthesis sites on DNA framework (Fig. 5b) with varying distances ranging from 110 nm to 10.9 nm and assembled TdT molecules at the corresponding sites. The successful assembly of TdT molecules at the designated sites and the ability to control the inter-site distances were confirmed via AFM imaging (Fig. 5c).

**Figure 5.**
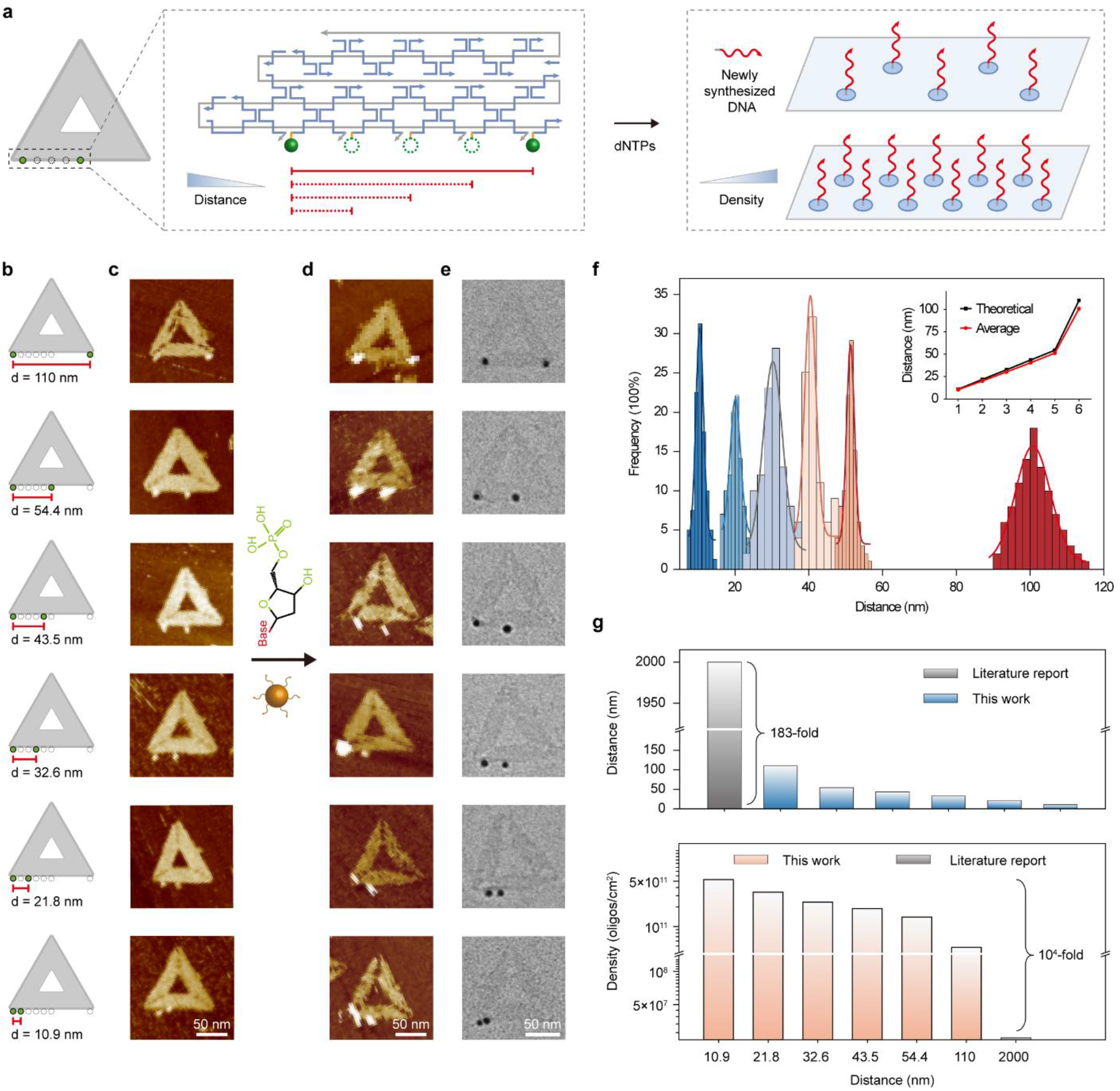
Distance regulation and DNA synthesis density control on DNA framework interface. (a) Schematic representation of distance regulation between synthetic sites and its impact on synthesis density on the DNA framework interface. DNA synthesis density increases as the distance between synthesis sites decreases. (b) Schematic of regulating the distance between adjacent synthetic sites on the DNA framework. (c) AFM imaging of TdT assembled on the DNA framework with varying distances between adjacent synthetic sites. Scale bar, 50 nm. (d) AFM imaging of newly synthesized DNA sequences with different distance on DNA framework by hybridized with SNAs. Scale bar, 50 nm. (e) TEM images of newly synthesized DNA on the DNA framework hybridized with SNAs. Scale bar, 50 nm. (f) Statistical analysis of distances between adjacent synthetic sites based on TEM images. (g) Calculation of theoretical synthesis density based on the distance between adjacent synthetic sites. As the distance decreases, the theoretical synthesis density increases significantly, with all values showing orders of magnitude higher than Microsoft’s DNA synthesis density.

Next, enzymatic DNA synthesis was performed, and the presence of newly synthesized sequences was confirmed by AFM and transmission electron microscopy (TEM) following SNAs hybridization (Figs. 5d, e). Statistical analysis of the distances between adjacent AuNPs revealed that the observed distances closely matched the theoretical values (Fig. 5f), demonstrating the precise programmability of our method. Compared to the 2 μm spacing between adjacent synthetic sites previously reported by Microsoft,^15^ our approach using DNA framework drastically reduces the distance to 10.9 nm, representing a 183-fold improvement. Accordingly, at this interval, we can achieve theoretical synthesis densities as high as 5.3 × 10 ^11^ oligonucleotides/cm^2^ (Fig. 5g), which is four orders of magnitude higher than Microsoft’s electrochemical synthesis method (2.5 × 10^7^ oligonucleotides/cm^2^). Furthermore, given that the single base synthesis rate of enzymatic DNA synthesis typically ranges from seconds to tens of seconds, our strategy could achieve a gigabyte per second (GB/s) data write rate for DNA data storage, a speed that is three orders of magnitude higher than Microsoft’s method. This significant improvement opens up new possibilities for high-speed, high-capacity data storage using DNA as the medium.

## Conclusions

In summary, we have developed an ultra-high throughput enzymatic DNA synthesis strategy with single-molecule-level precision based on DNA framework. By engineering TdT-DNA complexes on DNA frameworks, we precisely defined synthesis sites, leveraging the nanometer-scale addressability of DNA framework to control their position and spacing. This approach achieved a theoretical synthesis density of 5.3×10 ^11^ sequences per square centimeter, with adjacent synthesis sites spaced just 10.9 nm apart—representing a four-order-of-magnitude improvement over existing high-throughput synthesis platforms. Additionally, we demonstrated large-scale DNA synthesis on DNA framework arrays as well as the synthesis of unique DNA sequences to store “NGDS” information. While enzymatic DNA synthesis remains in its early stages and single-base precision remains a challenge, our work demonstrates a transformative pathway for scaling synthesis throughput and expanding the utility of DNA-based data storage. Further efforts, including the development of 3’ blocked nucleotides and engineered TdT with improved enzyme activity,^31^ will improve its application. Looking forward, we believe that harnessing multiplexed enzymatic DNA synthesis will usher in a new era of high-throughput DNA synthesis and digital data storage.

